# Constitutive discovery in the living human heart

**DOI:** 10.64898/2026.07.11.737831

**Authors:** Denisa Martonová, Fikunwa O. Kolawole, Sarthak A. Shinde, Daniel B. Ennis, Ellen Kuhl

## Abstract

Constitutive models of myocardial mechanics form a cornerstone of personalized cardiac simulations and cardiac digital twins. Researchers traditionally prescribe these models a priori and calibrate them from ex vivo tissue experiments, even though tissue excision alters loading conditions, removes residual stresses, and eliminates important physiological interactions. Multimodal cardiac MRI now provides subject-specific ventricular geometry, deformation, and myocardial microstructure, yet current inverse approaches still rely on predefined constitutive laws. Here we present the first framework to discover constitutive models of passive myocardial mechanics directly from in vivo cardiac imaging data by embedding a constitutive artificial neural network within a nonlinear finite element model of ventricular filling. Using multimodal cardiac MRI that combines ventricular geometry, deformation, and microstructure from a representative healthy individual, the framework identifies sparse, mechanically admissible strain-energy functions without prescribing their form a priori. The best-performing model contains only two fiber- and two sheet-invariant terms, achieves a mean displacement error of 1.62 mm, and reduces the error of the widely used Guccione and Holzapfel models by 34.14% and 26.01%. The discovered models indicate that fiber- and sheet-related anisotropic mechanisms dominate the passive mechanical response during physiological ventricular filling. More broadly, this work establishes a non-invasive strategy for subject-specific constitutive discovery from cardiac imaging data and lays the foundation for personalized cardiac simulations and cardiac digital twins.

## 1 Introduction

Cardiac mechanics play a central role in ventricular function and in the progression of cardiovascular disease. The passive mechanical response of the myocardium governs ventricular deformation during diastolic filling and determines the stress distribution within the ventricular wall. Alterations in myocardial stiffness associate with pathological conditions such as hypertrophy, fibrosis, and heart failure. Accurate constitutive models of myocardial tissue therefore form a cornerstone of computational cardiology, personalized heart simulations, and cardiac digital twins (Chabiniok et al. 2016; Peirlinck et al. 2021; Brown et al. 2026).

Numerous constitutive models have been proposed to describe the nonlinear mechanical behavior of soft biological tissues. Classical hyperelastic models, which include the neo-Hookean, Mooney-Rivlin, and Ogden models, express the strain-energy density as a function of invariants of the deformation tensor (Treloar 1944; Rivlin 1948; Ogden 1972). For anisotropic tissues, additional invariants account for interactions between deformation and embedded fiber families (Holzapfel et al. 2000). In cardiac biomechanics, structurally motivated constitutive laws such as the Guccione (GC) and Holzapfel-Ogden (HO) models explicitly represent myocardial microstructure and remain widely used in cardiac simulations (Guccione et al. 1991; Holzapfel and Ogden 2009). In practice, however, these constitutive models are prescribed *a priori*, and we need to identify their parameters from experimental or clinical data.

Traditionally, the constitutive behavior is characterized through ex vivo mechanical experiments on excised tissue specimens. Uniaxial and biaxial extension as well as shear tests provide detailed stress-strain relationships and reveal the anisotropic response of ventricular myocardium (Dokos et al. 2002; Sommer et al. 2015; Kakaletsis et al. 2021; Martonová et al. 2021). These experiments remain an important source of biomechanical insight and have enabled the calibration of increasingly sophisticated constitutive models. Nevertheless, ex vivo measurements alone cannot fully reproduce the physiological behavior of the beating heart. Tissue excision alters loading conditions, removes residual stresses, and eliminates interactions between myocardial tissue, blood pressure, and surrounding anatomical structures (Rodriguez et al. 1993; Grobbel et al. 2018).

To overcome these limitations, several studies have pursued inverse finite element (FE) approaches that estimate myocardial material parameters directly from in vivo measurements. Early work showed that tagged MRI measurements combined with nonlinear optimization and FE simulations can identify myocardial material parameters (Moulton et al. 1995, 1996). Subsequent studies incorporated myocardial fiber architecture obtained from cardiac diffusion tensor imaging (cDTI) (Augenstein et al. 2005, 2006) and investigated regional variations in myocardial stiffness associated with cardiovascular disease (Zhang et al. 2021). More recent efforts improved parameter estimation through advanced optimization strategies, combinations of ex vivo and limited in vivo data (Gao et al. 2015; Nasopoulou et al. 2017; Peirlinck et al. 2019; Lazarus et al. 2022; Laita et al. 2024) and the incorporation of heterogeneous full-field measurements (Krijnen et al. 2026). These studies demonstrated the potential of imaging-based and full-field inverse modeling. Recent studies have validated this approach using synthetic phantoms (Kolawole et al. 2023).

In parallel, advances in cardiac imaging have substantially expanded the biomechanical information available non-invasively in vivo. While earlier studies had to rely on a few surgically implanted videofluoroscopic markers to reconstruct strains, stresses, and stiffnesses in the beating heart (Krishnamurthy et al. 2008; Tsamis et al. 2011; Rausch et al. 2013), cine MRI now enables the reconstruction of exact ventricular geometry and motion, tagged MRI provides regional strain measurements, and cDTI characterizes myocardial microstructure. A recent multimodal dataset (Kolawole et al. 2025) combines all three modalities in the same individuals which provides a uniquely comprehensive description of cardiac anatomy, deformation, and fiber architecture. This rich dataset creates new opportunities for personalized constitutive characterization. Despite these advances, all inverse approaches remain restricted to predefined constitutive models whose particular form must be specified before parameter estimation.

Recent developments in material model discovery provide an alternative approach: Sparse regression methods demonstrated that governing equations can be identified directly from data without prescribing their functional form (Brunton et al. 2016). Building on these ideas, physicsinformed neural networks embed continuummechanics principles directly into network architectures while preserving material symmetries and thermodynamic consistency (Linka et al. 2021; Linka and Kuhl 2023; Linden et al. 2023). Combined with sparsification techniques such as *L*_1_ regularization, these approaches identify compact strain-energy functions directly from experimental observations (McCulloch et al. 2024). More recently, FE frameworks have allowed the direct learning of constitutive neural network models from heterogeneous full-field deformation data by integrating constitutive learning and FE simulation within a single optimization framework (Knipper et al. 2026). These developments extend constitutive learning far beyond traditional homogeneous stress-strain experiments.

Recently, automated constitutive discovery has been applied to myocardial tissue (Martonová et al. 2024; Martonová et al. 2025; Ingalkar et al. 2026). These studies showed that constitutive artificial neural networks (CANNs) identify compact strain-energy functions for human cardiac tissue from ex vivo mechanical experiments and suggested that only a small subset of invariant-based terms is sufficient to capture the anisotropic response of the myocardium. However, to date, these approaches rely exclusively on homogeneous laboratory experiments on excised tissue samples and do not exploit the rich biomechanical information available from multimodal in vivo cardiac imaging.

In this work, we extend constitutive model discovery from ex vivo tissue experiments to multimodal in vivo cardiac MRI. We combine cine MRI, tagged MRI, and cDTI to construct a personalized computational model that incorporates ventricular geometry, myocardial deformation, and myocardial fiber architecture. We formulate a general invariant-based strain-energy density function and use sparse regularization to identify the smallest set of constitutive terms that explains ventricular deformation during diastolic filling. Unlike conventional inverse FE methods that calibrate a predefined sparse constitutive model, our framework discovers the constitutive representation *directly* and entirely *autonomously* from in vivo observations. Here we focus on a representative healthy individual from a multimodal imaging cohort and systematically evaluate which invariant-based mechanisms the available imaging data can identify. This pilot study establishes a foundation for personalized constitutive model discovery from multimodal cardiac imaging and supports future developments in cardiac digital twins.

## 2 Methods

We build upon the multimodal cardiac MRI framework (Kolawole et al. 2025). This dataset combines cine MRI, tagged MRI, and cardiac diffusion tensor imaging (cDTI) acquired from healthy volunteers under a Stanford University Institutional Review Board-approved protocol with written informed consent. To isolate the influence of constitutive assumptions and model complexity, we focus on a single representative healthy volunteer.

Figure 1 schematically illustrates our workflow that combines multimodal cardiac MRI, finite element modeling, and automated constitutive model discovery. We reconstruct personalized ventricular geometry, deformation, and fiber architecture from imaging data and use these measurements to identify the constitutive model that best reproduces the observed ventricular deformation.

**Fig. 1.**
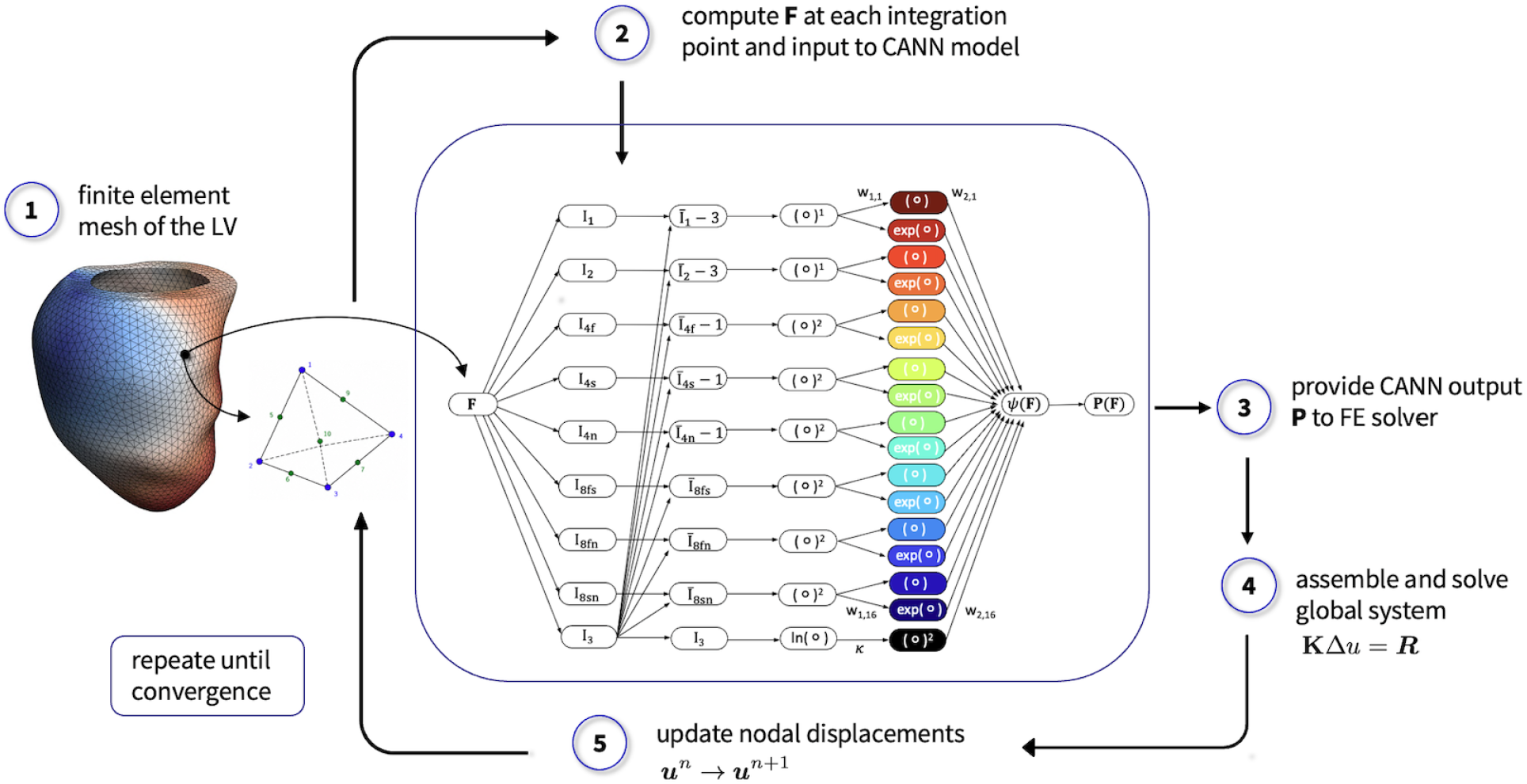
Computational workflow of the invariant-based CANN embedded within a finite element (FE) solver. It provides the deformation gradient **F** at each integration point from which the third and eight isochoric invariants are evaluated and passed through the CANN to obtain the strain-energy function *ψ* and the stress measure **P**. These quantities are used to assemble the global FE system, solve for the displacement increment, and update the configuration iteratively until convergence of both, the load increment and the loss in Eq. (12).

First, we construct a finite element model of the left ventricle from the cine MRI geometry. We use the diastasis configuration as the reference state and apply a physiological pressure load to the endocardial surface to reproduce ventricular filling up to end-diastole.

Next, we formulate a general strain-energy density function composed of invariant-based candidate terms. Sparse regularization selects the constitutive terms that best explain the data, while the optimization procedure identifies the corresponding material parameters by minimizing the discrepancy between simulated and MRI-derived displacement fields.

For the first time, our framework identifies both the constitutive model and its material parameters directly from in vivo measurements.

### 2.1 Continuum model

We describe the passive myocardium as a nearly incompressible hyperelastic material. The deformation mapping *φ* transforms a material point ***X*** in the reference configuration to its current position ***x***. The deformation gradient is

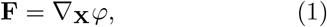

with determinant *J* = det(**F**). The right Cauchy– Green tensor is

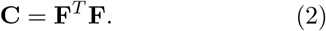

To separate volumetric and distortional deformations, we introduce the isochoric quantities

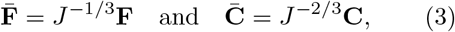

which satisfy 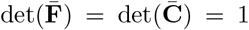. Hyperelastic materials derive their stress response from a strain-energy density function *ψ*, which we decompose into isochoric *ψ*_iso_ and volumetric *ψ*_vol_ parts,

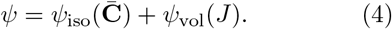

We obtain the first Piola-Kirchhoff stress as its derivative with respect to the deformation gradient,

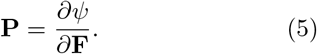

The myocardium exhibits pronounced anisotropy due to the organization of cardiomyocyte fibers into laminar sheet structures (Holzapfel and Ogden 2009; Sommer et al. 2015). Let **f**_0_, **s**_0_, and **n**_0_ denote the fiber, sheet, and sheet-normal directions in the reference configuration. We can express the constitutive response in terms of the volumetric invariant

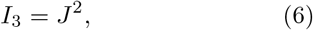

the isochoric isotropic invariants

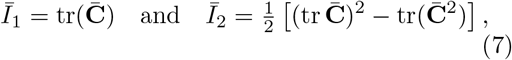

the isochoric anisotropic invariants

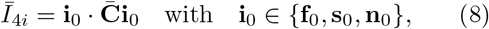

and the isochoric coupling invariants

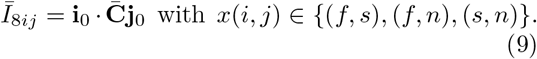

We obtain subject-specific fiber orientations from cDTI measurements acquired at end-systole and subsequently mapped to the diastatic reference configuration using the tagged-MRI-derived kinematic field (Kolawole et al. 2025). The resulting fiber field defines the local fiber direction **f**_0_. We define the normal direction **n**_0_ as the local transmural direction and compute the sheet direction **s**_0_ to fulfill the orthogonality condition, see Figure 2. This assumption is in agreement with experimental evidence that myocardial sheetlets lie approximately parallel to the wall in diastole (Nielles-Vallespin et al. 2017; Ghonim et al. 2017).

**Fig. 2.**
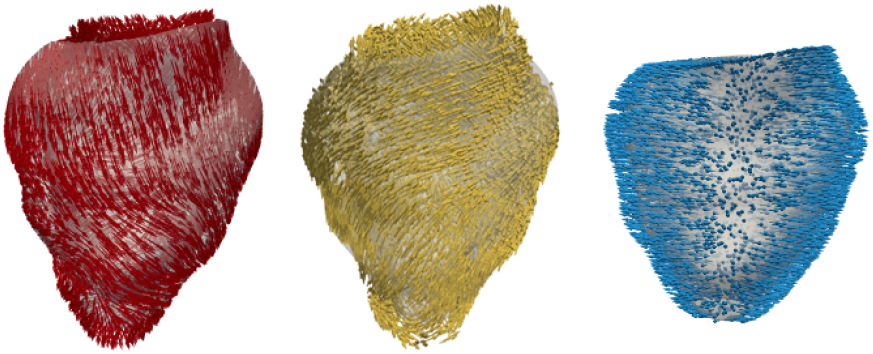
From left to right: Personalized fiber, sheet, and normal directions in the representative left ventricle.

### 2.2 Invariant-based constitutive model discovery

Classical myocardial constitutive models such as the transversely isotropic GC model (Guccione et al. 1991) and the invariant-based HO model (Holzapfel and Ogden 2009) can be viewed as particular combinations of constitutive features describing isotropic, fiber, and cross-fiber mechanical responses. Motivated by this observation and by recent data-driven model discoveries for myocardial tissue (Martonová et al. 2024; Ingalkar et al. 2026; Urrea–Quintero et al. 2026), we represent the strain-energy density using a constitutive neural network, displayed in the central part of the Figure 1, which operates directly on a set of isochoric invariants. The resulting strain-energy function is

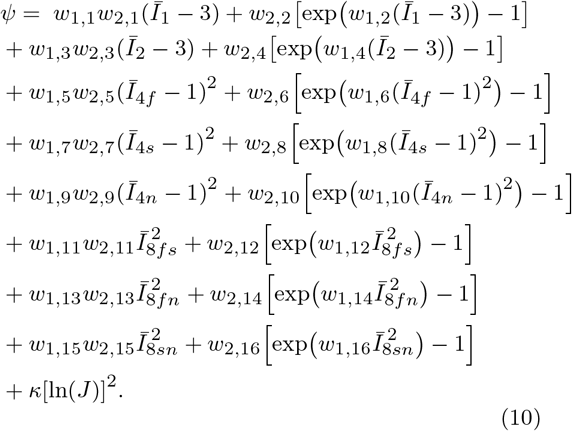

where the trainable parameters (*w*_*i,j*_) determine the contribution of the individual terms, while *κ* controls the volumetric penalty enforcing near incompressibility. The formulation combines isotropic, anisotropic, and interaction effects within a single constitutive representation and encompasses a broad class of hyperelastic models commonly used for soft biological tissues. The constitutive library, which consists of 2^16^ model combinations contains both polynomial and exponential functions of these invariants. The polynomial terms recover responses commonly used in Rivlin-type hyperelastic models (Mooney 1940; Rivlin 1948), whereas the exponential terms capture the nonlinear strain stiffening characteristic of biological tissues and form the basis of Holzapfel-Ogden type models (Holzapfel and Ogden 2009).

### 2.3 Unsupervised constitutive model discovery through finite element simulation

Conventional constitutive modeling is typically performed by fitting a predefined sparse constitutive model to ex vivo stress-strain measurements obtained from isolated tissue samples. In contrast, the present framework performs constitutive model discovery in vivo, directly at the organ scale. This implies that we embed the CANN model within a nonlinear FE model of passive left ventricular diastolic filling, and discover the constitutive model and parameters *directly* from observed myocardial deformation without requiring stress measurements.

We construct the FE model from the personalized left ventricular geometry at diastasis, which serves as the reference configuration. It contains 48,305 10-node quadratic tetrahedral elements and 74,933 nodes. Regional myocardial displacements between diastasis and end-diastole are obtained from tagged MRI and used as experimental observations. We simulate passive ventricular filling by prescribing an endocardial pressure of 1kPa, lying within the physiological range from 0.6 to 1.6 kPa reported for healthy subjects (Bouchard et al. 1971; Kolawole et al. 2025). We restrict the analysis to the filling phase between diastasis and end-diastole to avoid assumptions about the shape of the diastolic pressure waveform.

For a given parameter vector **w** = (*w*_1,1_, …, *w*_2,16_, *κ*), we evaluate the strain-energy density function of Eq. (10) at every integration point of the FE mesh. We compute the deformation gradient **F** and the associated invariant set from the current displacement field and supply them to the constitutive neural network. We then obtain the first Piola-Kirchhoff stress tensor according to Eq. (5) and the consistent material tangent

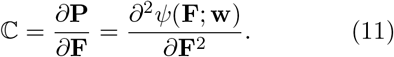

We assemble these quantities into the nonlinear finite element equilibrium equations and solve them iteratively using a Newton-Raphson procedure using the FEBio solver and user material plugin (Maas et al. 2012; Kakaletsis et al. 2021). We then identify the constitutive parameters by minimizing the error between simulated **u**_sim_ and MRI-derived **u**_MRI_ ventricular displacements. Specifically, we minimize the loss function

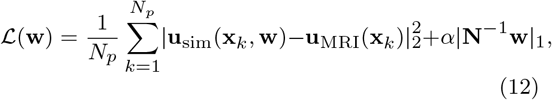

where the first term enforces agreement between simulated and observed ventricular kinematics, while the second term promotes sparse constitutive representations and improves parameter identifiability. Here *N*_*p*_ = 1084 denotes the number of myocardial mesh nodes located on seven equidistant horizontal planes (see Figure 3). We identify the mesh nodes closest to the tracked imaging planes and determine their positions at maximum inflation by interpolation of the three-dimensional tracking data, **N** is a normalization matrix to guaranty the equal contribution for all parameters and *α* controls the regularization strength. In this study, we set *α* = 0.01 (Martonová et al. 2025; Vervenne et al. 2025).

**Fig. 3.**
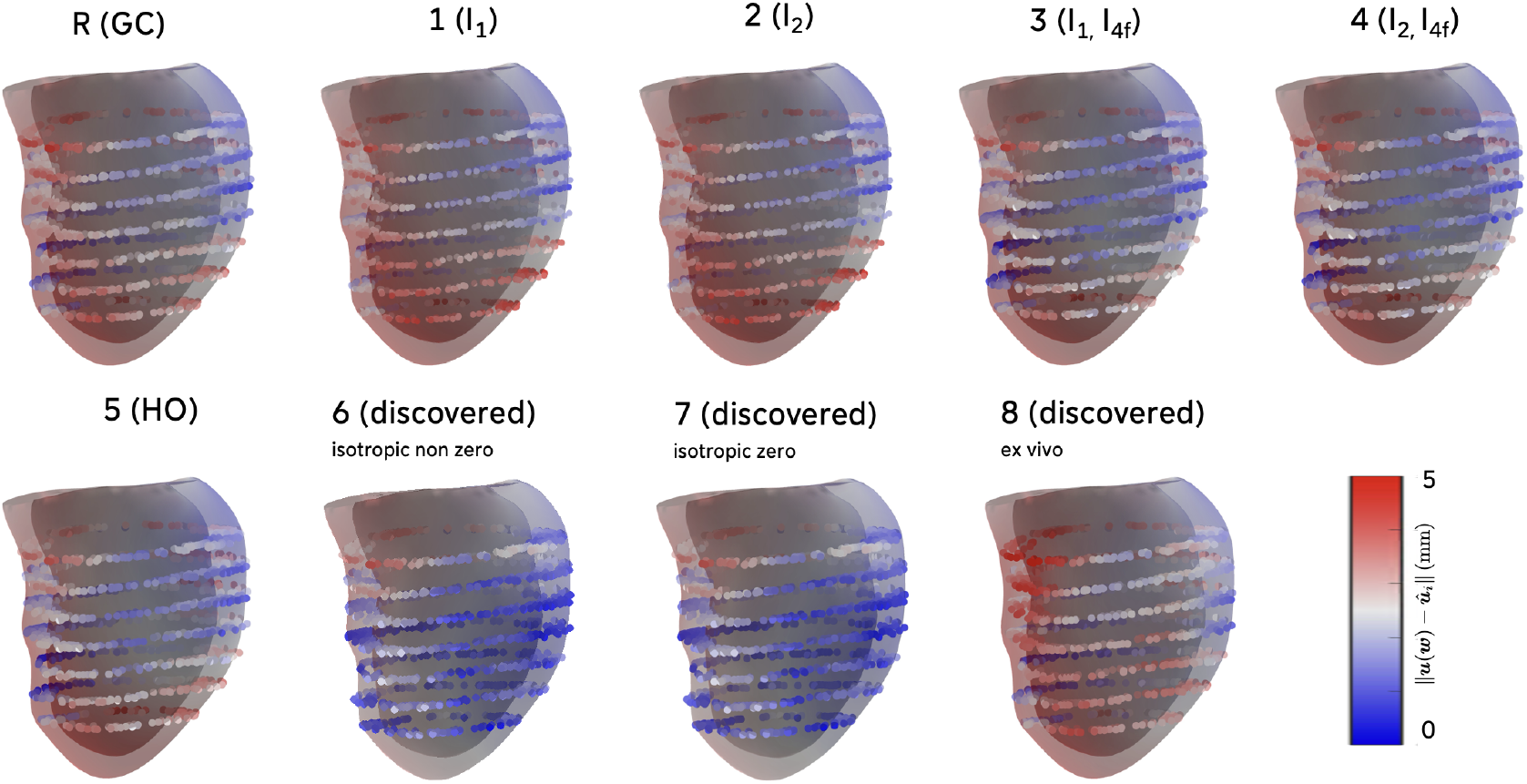
Displacement error between simulated and MRI-derived ventricular deformation at the end of the diastole. Corresponding prediction errors are displayed for the epicardial nodes used for the model discovery. Displayed left ventricular geometry is simulated with models and parameters specified in Tables 1 and 2.

For each optimization iteration, we evaluate the FE solution, compute the displacement mismatch with the MRI measurements, and determine how changes in the constitutive parameters affect this mismatch. We use these sensitivities to update the constitutive parameters and progressively reduce the loss function, see Figure 1.

**Table 1.** Displacement errors for the investigated constitutive models. Mean displacement error between simulated and MRI-derived ventricular deformation.

| model no. | active invariants | material symmetry / model | mean error [mm] | error change [%] |
| --- | --- | --- | --- | --- |
| R | – | transversely isotropic (Guccione) | 2.46 | 0.00 |
| 1 | $I_1$ | isotropic | 2.60 | +5.69 |
| 2 | $I_2$ | isotropic | 2.62 | +6.50 |
| 3 | $I_1, I_{4f}$ | transversely isotropic | 2.27 | -7.72 |
| 4 | $I_2, I_{4f}$ | transversely isotropic | 2.28 | -7.31 |
| 5 | $I_1, I_{4f}, I_{4s}, I_{8fs}$ | orthotropic (Holzapfel-Ogden) | 2.26 | -8.13 |
| 6 | $I_1, I_{4f}, I_{4s}, I_{8fn}, I_{8sn}$ | orthotropic (discovered, $I_1$ active) | 1.71 | -30.49 |
| 7 | $I_{4f}, I_{4s}, I_{8fn}, I_{8sn}$ | orthotropic (discovered) | 1.62 | -34.14 |
| 8 | $I_2, I_{4f}, I_{4n}, I_{8fs}$ | orthotropic (discovered ex vivo) | 2.70 | +9.76 |

**Table 2.** Strain-energy functions and parameters. Non-zero terms of the predefiened (1-5) and discovered (6-7) constitutive models. We report parameters as *w*_1,*•*_ (dimensionless), *w*_2,*•*_ (kPa) and *κ* (MPa), see Eq. (10).

| network weights | model term | model number according to Tab. 1 |  |  |  |  |  |  |
| --- | --- | --- | --- | --- | --- | --- | --- | --- |
|  |  | 1 | 2 | 3 | 4 | 5 | 6 | 7 |
| $w_{1,1} \cdot w_{2,1}$ | $\bar{I}_1 - 3$ | 12.09 | — | — | — | — | 1.00, 0.50 | — |
| $w_{1,3} \cdot w_{2,3}$ | $\bar{I}_2 - 3$ | — | 12.92 | — | — | — | — | — |
| $w_{1,2}, w_{2,2}$ | $\exp(\bar{I}_1 - 3)$ | — | — | 8.75, 0.56 | — | 0.98, 5.39 | — | — |
| $w_{1,4}, w_{2,4}$ | $\exp(\bar{I}_2 - 3)$ | — | — | — | 13.87, 0.31 | — | — | — |
| $w_{1,6}, w_{2,6}$ | $\exp[(\bar{I}_{4f} - 1)^2]$ | — | — | 10.51, 1.65 | 8.93, 1.93 | 100.00, 0.12 | 38.64, 0.72 | 38.78, 0.77 |
| $w_{1,8}, w_{2,8}$ | $\exp[(\bar{I}_{4s} - 1)^2]$ | — | — | — | — | 90.76, 0.003 | 47.64, 0.03 | 48.71, 0.04 |
| $w_{1,12}, w_{2,12}$ | $\exp(\bar{I}_{8fs}^2)$ | — | — | — | — | 9.42, 0.75 | — | — |
| $w_{1,14}, w_{2,14}$ | $\exp(\bar{I}_{8fn}^2)$ | — | — | — | — | — | 8.23, 2.20 | 8.34, 2.03 |
| $w_{1,16}, w_{2,16}$ | $\exp(\bar{I}_{8sn}^2)$ | — | — | — | — | — | 8.30, 2.07 | 8.36, 1.97 |
| $\kappa$ | $[\ln(J)]^2$ | 2.55 | 2.55 | 2.55 | 2.55 | 2.55 | 2.85 | 2.96 |

### 2.4 Systematic investigation of activated invariants and model complexity

To investigate the influence of constitutive assumptions on the myocardial response, we perform a sequence of constitutive discovery studies with progressively increasing model complexity: In the first stage, we restrict the constitutive model to the isotropic invariants *Ī*_1_ and *Ī*_2_ which yields a purely isotropic hyperelastic material model. In the second stage, we additionally include the fiber invariant *Ī*_4*f*_ to account for fiber-direction anisotropy and obtain a transversely isotropic constitutive response. In the third stage, we restrict the constitutive model to the invariant set underlying the classical HO model (Holzapfel and Ogden 2009) which enables a direct comparison with a widely used constitutive model of passive myocardium. In the final stage, we perform unrestricted constitutive discovery and use the complete invariant-based strain-energy function defined in Eq. (10). We allow the CANN to utilize all isotropic, anisotropic, and interaction invariants as well as the bulk modulus *κ*, and optimize all constitutive parameters simultaneously.

In contrast to classical constitutive formulations, which prescribe the relative importance of fiber, sheet, sheet-normal, and coupling contributions *a priori*, the proposed framework determines these contributions directly from the imaging data. At the same time, the discovery process remains physically constrained as the constitutive response is restricted to the invariant-based strain-energy representation of Eq. (10) and embedded within the FE model. Furthermore, sparsity-promoting regularization suppresses inactive constitutive mechanisms, limits model complexity, and reduces the risk of overfitting. The resulting framework therefore avoids both the structural assumptions of predefined constitutive laws and the unconstrained nature of black-box neural-network models.

Throughout all studies, we enforce positivity constraints on the trainable parameters to ensure non-negative strain-energy contributions and a physically admissible material behavior. Based on the previous studies, we also set an upper parameter limit of 100 kPa on the stress-like weights to avoid non-physiological values (Guan et al. 2019; Martonová et al. 2024).

The hierarchical investigation with increasing model complexity allows us to quantify the influence of anisotropy, invariant selection, and constitutive complexity on the discovered strain-energy function and the resulting prediction of ventricular mechanics.

## 3 Results

Table 1 summarizes the displacement errors for all investigated constitutive models, while Figure 3 shows the corresponding spatial distribution of displacement mismatches between simulated and MRI-derived ventricular kinematics. The discovered models and parameters for the specific cases are summarized in Table 2.

The transversely isotropic Guccione model calibrated through a single stiffness parameter (Kolawole et al. 2025) yields a mean displacement error of 2.46 mm and serves as a reference. Restricting constitutive discovery to isotropic constitutive responses based on either *Ī*_1_ or *Ī*_2_ increases the displacement error, which indicates that isotropic constitutive descriptions cannot adequately reproduce the measured ventricular deformation. When we include the fiber invariant *Ī*_4*f*_, as expected, the predictive accuracy reduces the error below the Guccione baseline. This highlights the importance of myocardial anisotropy.

Among the predefined constitutive structures, the orthotropic Holzapfel-Ogden model achieves the lowest mean displacement error of 2.26 mm. Constitutive discovery identifies several orthotropic invariant combinations that further improve agreement with the imaging data. The best-performing model activates the invariants *Ī*_4*f*_, *Ī*_4*s*_, *Ī*_8*f n*_, *Ī*_8*sn*_, and reduces the mean displacement error to 1.62 mm which corresponds to a reduction of 34.1% relative to the Guccione model. Figure 3 shows that the ventricular wall is dominated by blue-colored low-error regions for this constitutive model.

The choice of constitutive model influences not only the displacement prediction error at end-diastole but also the resulting stress distribution. As illustrated in Figure 4, different models lead to distinct stress profiles. Pronounced differences are observed in the von Mises stress field, particularly in regions of high stretch located in the lower part of the left ventricle. For the anisotropic models (R, 3–7), the von Mises stress reaches values of approximately 5 kPa in these regions. In contrast, the isotropic models (1, 2) produce a more homogeneous stress distribution with less pronounced stress concentrations.

**Fig. 4.**
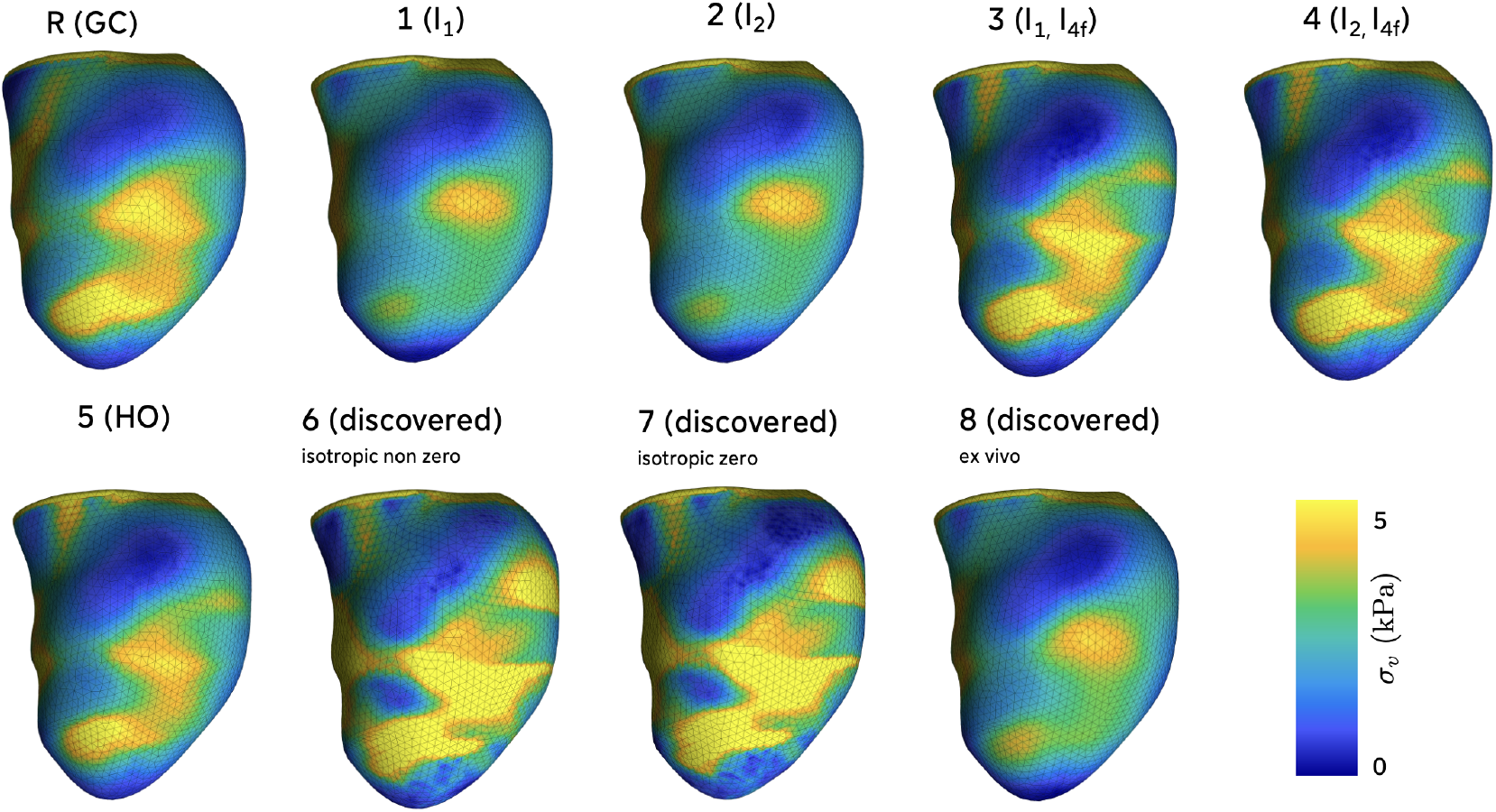
Stress profiles across human left ventricle at the end of the diastole. Von Misses stress generated by the left ventricle is simulated with models and parameters specified in Tables 1 and 2.

To assess generalization beyond the ventricular fitting problem, we evaluate the performance of the discovered constitutive model under ex vivo loading protocols (Sommer et al. 2015; Martonová et al. 2024). While the in vivo-derived constitutive models reproduce the ventricular deformation with high accuracy, none of them reproduces the ex vivo stress-stretch relationships. Figures 5 and 6 illustrate that all loading curves yield a mismatch between predicted and experimental data with a negative goodness of fit, *R*^2^ *<* 0. In contrast, the constitutive model and parameters previously identified from the ex vivo experiments (Sommer et al. 2015) also perform poorly when we simulate diastolic filling and yield the largest mean displacement error of 2.70 mm among all anisotropic models, see Table 1. Taken together, these observations emphasize an important finding: Constitutive descriptions that best explain ex vivo tissue mechanics do not necessarily reproduce in vivo ventricular deformation, and vice versa.

**Fig. 5.**
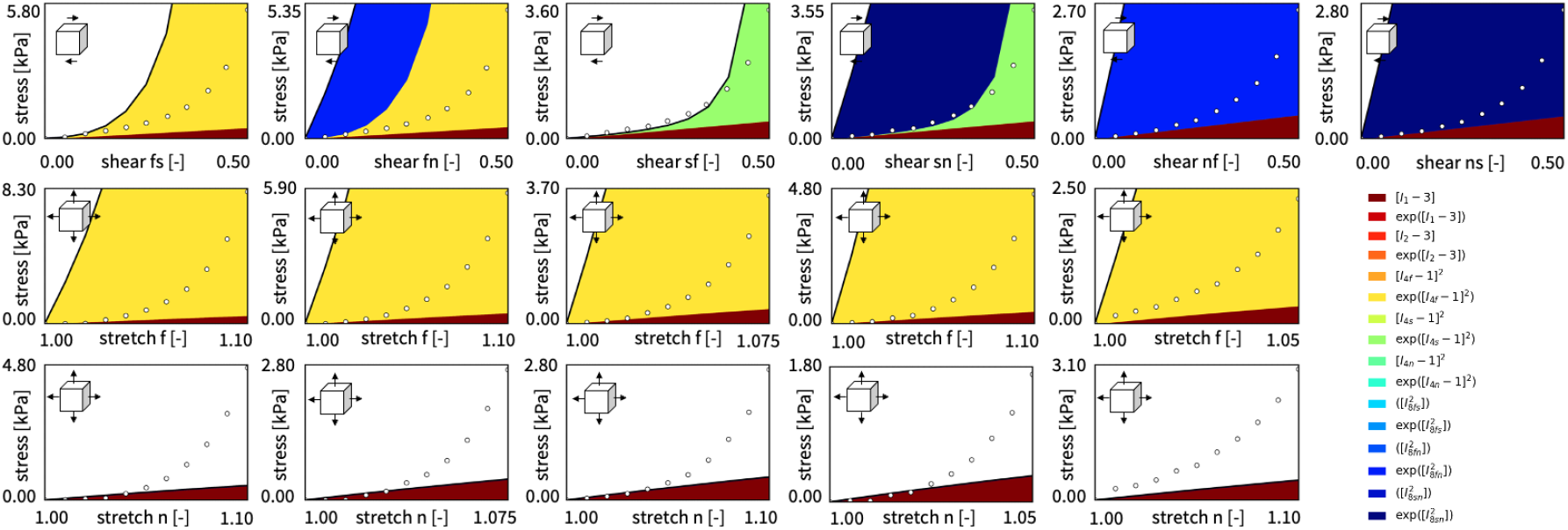
Comparison of constitutive responses by the discovered model under ex vivo loading protocols (six simple shear and five biaxial tests). Colored regions indicate the contributions of the active constitutive terms and symbols denote the experimental data (Sommer et al. 2015).

**Fig. 6.**
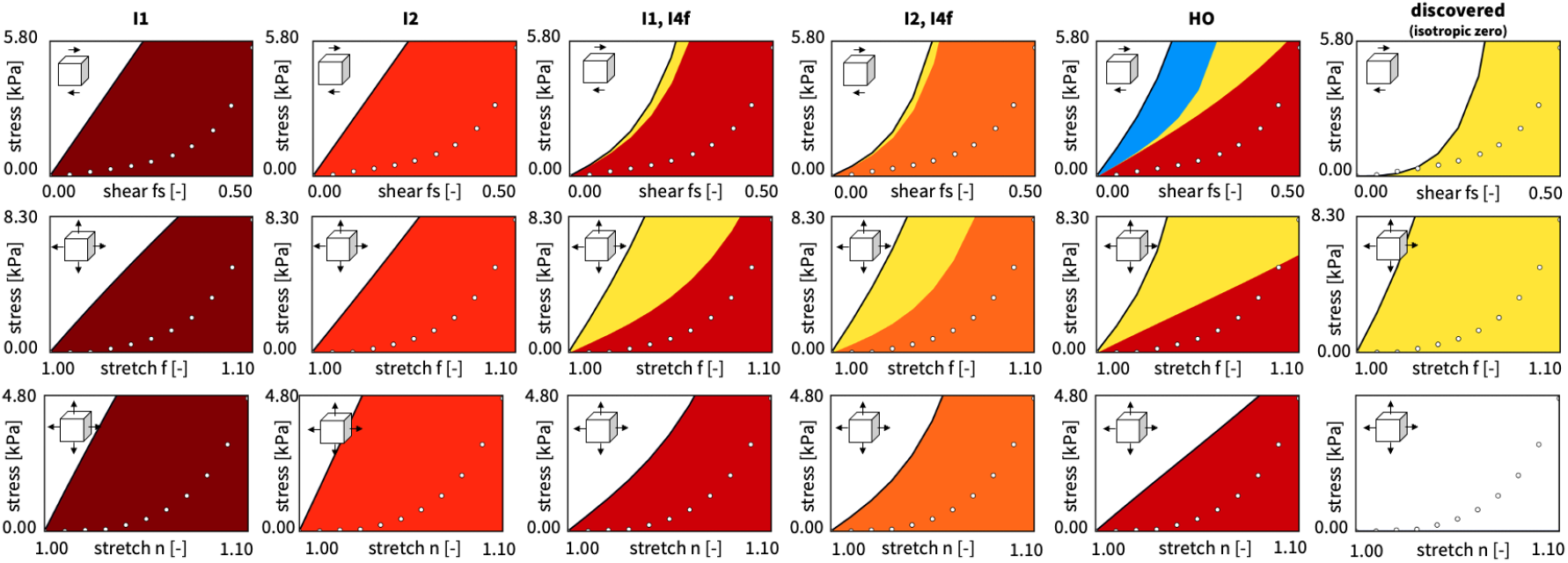
Comparison of different constitutive model responses under ex vivo loading protocols (one simple shear and one biaxial tests). Colored regions show the contributions of the active constitutive invariants and dots indicate experimental data (Sommer et al. 2015).

## 4 Discussion

The present pilot study demonstrates that constitutive model discovery from multimodal in vivo cardiac MRI identifies constitutive models that outperform established myocardial constitutive models in reproducing ventricular deformation at the end-diastole. At the same time, the constitutive models identified from in vivo ventricular deformation are not consistent with those discovered from independent ex vivo mechanical experiments. Together, these findings provide insight into the constitutive information encoded in ventricular filling data and reveal fundamental differences between the constitutive behavior identified at the organ scale and the constitutive behavior measured in isolated tissue samples.

### Ventricular filling activates distinct constitutive mechanisms

Our results showcase the dominant role of myocardial anisotropy in reproducing ventricular deformation. Restricting constitutive discovery to isotropic models based on either *Ī*_1_ or *Ī*_2_ increases the displacement error relative to the Guccione model, whereas inclusion of the fiber invariant *I*_4*f*_ substantially improves predictive accuracy. This observation agrees with extensive experimental evidence that passive myocardial mechanics are governed primarily by an anisotropic microstructural architecture and cardiomyocyte alignment (Guccione et al. 1991; Holzapfel and Ogden 2009; Sommer et al. 2015).

Interestingly, the isotropic invariants *Ī*_1_ and *Ī*_2_ provide nearly identical predictive capability. Models based solely on *Ī*_1_ or *Ī*_2_ produce comparable displacement errors and stress field, and this observation remains unchanged after introducing fiber anisotropy through *Ī*_4*f*_ . This result contrasts with previous constitutive discovery studies based on ex vivo mechanical experiments, where *Ī*_2_ often emerges as the dominant isotropic invariant while *Ī*_1_ contributes little additional predictive value (Linka et al. 2023; Kuhl and Goriely 2024; Martonová et al. 2026). The present findings suggest that end-diastolic ventricular filling does not sufficiently excite the deformation modes required to distinguish between competing isotropic constitutive mechanisms.

The best-performing constitutive models consistently include the fiber invariant *Ī*_4*f*_ together with sheet- and coupling invariants such as *Ī*_4*s*_, *Ī*_8*f n*_, and *Ī*_8*sn*_. Moreover, the discovery procedure suppresses many candidate constitutive terms. The resulting sparse constitutive model therefore suggests that physiological ventricular filling activates only a subset of the constitutive mechanisms commonly assumed in myocardial constitutive models.

### In vivo and ex vivo discovery yield different constitutive models

The ex vivo experiments in Figures 5 and 6 provide additional insight into the constitutive mechanisms identified from ventricular deformation. The discovered response is dominated by anisotropic contributions associated with the fiber and sheet directions, primarily through *Ī*_4*f*_ and *Ī*_4*s*_. Consistent with this observation, the sparse model with *Ī*_4*f*_, *Ī*_4*s*_, *Ī*_8*f n*_, *Ī*_8*sn*_ achieves the lowest ventricular displacement error even without featuring either of the isotropic invariants *Ī*_1_ or *Ī*_2_ or the normal invariant *Ī*_4*n*_. This constitutive model explains many aspects of the ex vivo predictions. The model reproduces substantial stiffness in loading modes associated with the fiber and sheet directions but exhibits zero resistance in deformation modes dominated by the normal direction. Since neither *Ī*_1_ nor *Ī*_2_, nor *Ī*_4*n*_ contribute significantly to the strain-energy density, the model contains no direct constitutive mechanism that generates stiffness under pure normal stretch.

Although the discovered constitutive models do not reproduce the ex vivo stress-stretch curves quantitatively, they still capture the qualitative anisotropic trends that we observe experimentally. The fiber-related loading modes generally exhibit the largest stiffness, whereas loading modes dominated by the cross-fiber directions remain substantially softer. At the same time, all loading curves yield negative coefficients of determination, *R*^2^ *<* 0, which indicates that none of the in vivo-derived constitutive models reproduce the ex vivo measurements with meaningful quantitative accuracy.

A similarly interesting observation emerges in the opposite direction. The constitutive model and parameters previously identified from the ex vivo experiments (Sommer et al. 2015) yield the largest displacement error among all investigated anisotropic constitutive models when evaluated in the ventricular fitting problem. Thus, the constitutive description that best explains the ex vivo experiments does not provide the best description of ventricular deformation, while the constitutive model that best reproduces ventricular deformation does not reproduce the ex vivo stress-stretch data. This agrees with recently reported discrepancies between ex vivo and in vivo constitutive models (Peirlinck et al. 2019; Lazarus et al. 2022; Laita et al. 2024).

Several factors may contribute to these differences: First, the present study identifies constitutive behavior from a personalized left-ventricular geometry, personalized fiber architecture, and personalized deformation measurements obtained from a single healthy volunteer. In contrast, the ex vivo experiments (Sommer et al. 2015) represent average tissue responses obtained from multiple myocardial specimens and donors. Consequently, part of the observed discrepancy may reflect biological variability rather than differences in constitutive structure alone.

Second, in vivo constitutive identification reflects the effective mechanical response required to reproduce ventricular deformation under physiological loading conditions. The identified constitutive parameters therefore implicitly account for the influence of ventricular geometry, heterogeneous fiber architecture, residual stress, ventricular tethering, and tissue interactions that are absent in isolated tissue specimens (Peirlinck et al. 2019; Kolawole et al. 2025). Ex vivo experiments remove many of these effects and instead characterize the response of small tissue samples under highly controlled loading conditions. The resulting constitutive parameters therefore represent different effective descriptions of myocardial mechanics.

Third, the constitutive model identified in the present study depends on assumptions regarding ventricular loading and boundary conditions. The inverse problem relies on prescribed end-diastolic pressure and kinematic constraints at the ventricular base, both of which influence the stress field required to reproduce the observed deformation. Uncertainties in pressure measurements, reference-state definition, or boundary conditions may therefore affect the constitutive parameters identified from the imaging data. Since the present framework uses a single loading state corresponding to end-diastolic filling, it remains difficult to fully separate constitutive effects from uncertainties in loading conditions.

### The normal direction remains weakly constrained during filling

The interpretation of the weak *Ī*_4*n*_ contribution requires additional care when comparing the present results with ex vivo experiments. In vivo studies of the beating heart have shown that myocardial sheetlets are arranged nearly parallel to the ventricular wall during diastole and reorient toward a more transmural configuration during systole (Nielles-Vallespin et al. 2017; Ghonim et al. 2017). More recently, nested-tori representations have been proposed to capture this ventricular microstructure and its organization more accurately (Osouli et al. 2025). However, the mechanical state represented by ex vivo tissue is often uncertain, which complicates the definition of cross-fiber directions.

As a result, no consensus exists on how to define sheet and normal directions in ex vivo myocardial samples. Many studies assume that the sheet direction **s**_0_ is aligned with the transmural direction (Göktepe et al. 2011; Guan et al. 2019; Holz et al. 2023), while other studies define the normal direction **n**_0_ as transmural (Vaverka and Bursa 2024; Kakaletsis et al. 2021). Consequently, the interpretation of sheet- and normal-related constitutive mechanisms should account for potential differences in how cross-fiber directions are defined and represented.

To further investigate this effect, we additionally constrained the isotropic contribution to remain active by enforcing non-zero parameters associated with the isotropic invariant *Ī*_1_. This modification restored non-zero stresses in loading modes that were previously inactive while preserving the overall anisotropic response. The resulting model remained dominated by fiber- and sheet-related mechanisms and indicates that ventricular filling strongly constrains these constitutive directions, while the normal direction remains only weakly constrained.

### Model discovery supports a near-incompressible behavior

The constitutive discovery framework identifies a volumetric penalty parameter of *κ* ≈ 2.9 MPa in the logarithmic volumetric energy term Ψ_vol_ = *κ* [ln(*J* )]^2^. This parameter penalizes deviations from *J* = 1 and therefore promotes near-incompressible behavior during ventricular filling. Model discovery consistently identifies a finite value of *κ* rather than driving the material toward strict incompressibility. This result agrees with experimental observations that perfused myocardial tissue undergoes small but measurable volume changes and therefore does not satisfy exact incompressibility (Bonnemains et al. 2019). At the same time, the magnitude of the identified parameter remains sufficiently large to suppress substantial volume changes, which suggests that the in vivo response lies close to the near-incompressible limit. Consequently, the *I*_3_-dependent term plays an important role in enforcing physiological volume preservation, even though the data do not support an exactly incompressible constitutive response.

### Constitutive discovery remains mechanically admissible by construction

The proposed framework shares the objective of learning constitutive behavior from experimental observations with physics-informed neural networks (PINNs) (Raissi et al. 2019). However, the underlying methodology differs fundamentally: Conventional PINN approaches typically approximate displacement fields using neural networks and enforce equilibrium, constitutive equations, and boundary conditions through additional residual terms in the loss function. Consequently, the optimization requires balancing multiple competing objectives via tunable weighting factors. As a result, the final solution is *highly sensitive* to user-defined weights.

In contrast, the present approach embeds the constitutive artificial neural network (CANN) directly within a nonlinear finite element solver. Mechanical equilibrium, compatibility, and boundary conditions are therefore satisfied by construction throughout the optimization. Rather than learning a displacement field, we learn the constitutive law itself and directly evaluate its goodness of fit through the mismatch between simulated and measured ventricular deformation. This formulation eliminates the need for balancing displacement, constitutive, and equilibrium residuals and ensures that every constitutive update *remains mechanically admissible*.

### Implications, limitations, and future directions

The present findings have important implications for constitutive model discovery from medical imaging data. Importantly, we should not interpret the discovered model as the single unique constitutive law for myocardial tissue. Instead, it represents the constitutive model that *best explains the available in vivo imaging data*. Different loading conditions may very well activate additional constitutive mechanisms and therefore lead to different constitutive models. Yet, to which extent these alternative loading conditions are relevant under physiological conditions remains to be more thoroughly explored.

This perspective aligns with recent developments in constitutive model discovery (Linka et al. 2023; McCulloch et al. 2024; Knipper et al. 2026).

Rather than assuming a constitutive law *a priori*, these approaches seek the simplest constitutive representation supported by the available data. The present study demonstrates that this concept can be extended from ex vivo mechanical experiments to multimodal in vivo cardiac MRI. The current study focuses on a single healthy volunteer and a single loading state corresponding to passive ventricular filling. This setting provides a controlled environment for investigating constitutive discovery, but may inevitably limit generalizability. Additional loading states, multiple cardiac phases, pressure perturbations, or larger imaging cohorts may activate constitutive mechanisms that remain weakly constrained in the present framework. Future studies should also investigate richer libraries of constitutive features and extend the framework to active contraction and electromechanical coupling.

Despite these limitations, the proposed framework demonstrates that constitutive model discovery can be performed non-invasively using multimodal in vivo cardiac MRI, while preserving mechanical consistency through the finite element formulation. The results suggest that ventricular deformation contains sufficient information to identify a sparse set of dominant constitutive mechanisms, but not necessarily enough information to uniquely recover tissue-scale constitutive behavior. These findings provide a foundation for future personalized constitutive modeling and cardiac digital twins (Peirlinck et al. 2021).

## 5 Conclusion

We present the first framework for constitutive model discovery directly from in vivo cardiac imaging data by embedding a constitutive artificial neural network within a non-linear finite element model of ventricular filling. Using subject-specific ventricular geometries, cDTI-derived microstructures, and MRI-derived displacement measurements, the framework identifies constitutive representations without the need to prescribe a constitutive law a priori. In contrast to physics-informed neural networks, this approach eliminates the need for balancing displacement, constitutive, and equilibrium residuals and ensures that every constitutive update remains mechanically admissible. It discovers constitutive models directly from cardiac imaging data that consistently outperform the widely used Guccione and Holzapfel-Ogden models during ventricular filling. Notably, the best-performing sparse model, with only two fiber- and two sheet-invariant terms, achieves a mean displacement error of 1.62 mm and reduces the error of the Guccione and Holzapfel models by 34.14% and 26.01%. Beyond improved predictive accuracy, the discovered models indicate that fiber- and sheet-related anisotropic mechanisms dominate the passive mechanical response during physiological ventricular filling. Taken together, the proposed approach provides a mechanically consistent strategy for discovering subject-specific constitutive models non-invasively and directly from cardiac imaging data and establishes a foundation for personalized cardiac simulations and digital twin modeling.

## Supplementary information

Our source code, data, and examples are available at https://github.com/LivingMatterLab/CANN.

## Acknowledgments

The authors acknowledge support from the European Research Council (ERC) Grant 101141626 DISCOVER funded by the European Union to EK; views and opinions expressed are, however, those of the authors only and do not necessarily reflect those of the European Union or the ERC Executive Agency. Neither the European Union nor the granting authority can be held responsible for them; from FAU Emerging Talents Initiative (ETI) to DM and from NIH R01 HL131823 to DBE.

## Notes

### Competing Interest Statement

The authors have declared no competing interest.

